# Mutant glucocorticoid receptor binding elements on Interleukin-6 promoter regulate dexamethasone effects

**DOI:** 10.1101/2020.06.24.168666

**Authors:** Wen-Teng Chang, Ming-Yuan Hong, Chien-Liang Chen, Cheng-Chieh Tsai, Chi-Yuan Hwang, Chia-Chang Chuang

**Affiliations:** Department of Biological Science and Technology, Chung Hwa University of Medical Technology, Tainan, Taiwan; Department of Emergency Medicine, National Cheng Kung University Hospital, College of Medicine, National Cheng Kung University, Tainan, Taiwan; Department of Physical Therapy, I-Shou University, Kaohsiung, Taiwan; Department of Nursing, Chung Hwa University of Medical Technology, Tainan 701, Taiwan

**Keywords:** Glucocorticoid receptor, interleukin-6, lipopolysaccharide, dexamethasone, promoter activity, transcriptional factor

## Abstract

Glucocorticoid has been widely used as an important modulator for clinical infectious and inflammatory disease. Glucocorticoid receptor (GR) is a transcription factor belonging to the family of nuclear receptors, regulated anti-inflammatory process and the release of pro-inflammatory cytokines. Five putative GR and other transcription factor binding sites on interleukin (IL)-6 promoter were identified and dexamethasone could reduce LPS-induced IL-6 release. Among them, the mutant transcriptional factors NF-κB, AP-1, and Sp1-2 site decreased the basal and effects of lipopolysaccharide (LPS)-induced IL-6 promoter activities in different responses. GR2/3 seemed to be an important role in both basal and inducible promoter activities in LPS-induced inflammation. We concluded that the selective GR2/3 modulators may have agonistic and antagonistic combined effects and activate important signaling pathway during LPS-stimulated inflammatory process.

## Introduction

Severe sepsis has been assumed to relate with immune dis-equilibrium. The immune dysregulation in the early phase of sepsis is result from inadequate endogenous glucocorticoid mediated regulation of nuclear factor-κB (NF-κB) activation, which leads to its over-expression and release massive pro-inflammatory cytokines (the so-called “cytokine storm”) [1]. “Glucocorticoid sensitivity” in critically ill patient with septic shock is also associated with disease severity and outcome [2]. Administration of low low-dose steroid in the early septic phase, although could not reduce 28-day mortality, it could reduce duration of shock and mechanical ventilator-dependent [3–5]. The effect of low-dose steroid in short- and longer mortality of septic patients remain controversial. Glucocorticoids (GCs) regulate many biological processes through their intracellular glucocorticoid receptors [GRs]. The GCs diffuse through the cell membrane bind to its GRs and the activated GRs complex translocated into the nucleus and expressed inflammatory mediators [6].

GCs are considered as immunosuppressive and anti-inflammatory agents and adjuvant therapy for patients with severe sepsis and septic shock. The activated GR is proposed to directly and indirectly interact with promoters to perform the negative regulation of cytokine gene expression [7,8]. Selective knockout of GR gene results in sensitive to LPS treatment, whereas, endogenous GCs decrease LPS-induced inflammation via inhibition of cytokine gene expression, such as IL-12 [9]. GCs also negatively regulate IL-6 gene expression through downregulation of its promoter activity in various tissues [10,11]. The regulation of several cytokine gene expressions by activated GRs, which are interact with the GCs, may be through binding to genes or indirective interaction with other transcription factors, in particular NF-κB and activator protein-1 (AP-1) complexes [7,12–14]. Transcription factors, such as NF-κB and AP-1, are known to mediate gene expression during inflammatory reaction extensively, especially in inflammatory cytokines, MMPs and cyclo-oxygenase-2 [12,15,16]. The synthetic glucocorticoid, dexamethasone (DEX), has been shown to inhibit inflammatory response via downregulation of transcription factor AP-1 in human lung epithelial cells [16].

We have previously demonstrated that the increase of macrophage migration inhibitory factor (MIF) was noted in severe sepsis such as *Vibrio vulnificus-*infected model [17]. In *V. vulnificus*-infected mice, MIF could regulate interleukin (IL)-6 in a time-dependent manner. Serum MIF regulates the NF-κB to modulate the release of IL-6 in transcriptional level. We further to investigate the mechanisms of transcription factors involved in regulating IL-6 promoter. We attempt to elucidate the roles of putative transcription factor binding sites, such as NF-κB, Sp1 and AP1, play in LPS-induced sepsis model. We also intend to depict the detailed causal link between GRs and pro-inflammatory transcription factors.

## Materials and Methods

### Cell Culture

Two cell lines were used in our studies. One was the RAW264.7 mouse macrophage cell lines, purchased from Culture Collection and Research Center, Food Industry and Development Institute, Hsinchu, Taiwan. Cells were grown in Dulbecco’s Modified Eagle Medium (Sigma) supplemented with 10% fetal bovine serum (Gibco Laboratories, Grand Island, NY) in a humidified atmosphere containing 5% CO2 at 37 °C. The second cell line was human neuroblastoma IMR-32 cells, obtained from the American Type Culture Collection (ATCC), were cultured in minimum essential medium (MEM) (Genedire X, USA) supplemented with 10% fetal bovine serum and 1% sodium pyruvate at 37°C in a controlled of 5% CO2 atmosphere. After 2-to 3-day growth, up to 70-80% full of 10 cm culture dish, the IMR-32 cultures were subcultured or collected and extracted out the nuclear proteins for EMSA use.

### Chemicals

Lipopolysaccharide (Escherichia coli O111: B4) and dexamethasone (DEX) were purchased from Calbiochem® (Detroit, MI) and Sigma Chemical Co. (St. Louis, MO), respectively.

### RNA isolation and semi-quantitative RT-PCR analysis

Total RNA was isolated from the cultured cells using TRIzol reagent (Invitrogen). RT-PCR was performed as described previously (Chang and Huang, 2005). Briefly, total RNA (2 μg) was reverse-transcribed into cDNA in 20 μl of 1X first strand buffer containing 0.5 μg of oligo(dT) as a primer, 500 μM dNTP, and 200 units of SuperScript II (Invitrogen). PCR was performed in 20 μl of 1X PCR buffer containing 2 μl of RT products, 1 unit of AmpliTaq DNA polymerase (Roche Applied Science), 200 μM dNTP, and 1.5 mM MgCl_2_ (Amersham Biosciences), and 0.4 μM primer pair. We used the primer pair of IL-6 mRNA, mIL6-mF1 and mIL6-mR1 (shown in Supplemental information). The PCR parameters were 94 °C for 30 s, 53 °C for 30 s, and 72 °C for 30 s for 30 cycles, followed by a final elongation at 72 °C for 7 min. PCR products were analyzed on a 1.2% agarose gel.

### Plasmid constructs

pGL4.10-Basic and pRL-TK luciferase reporter vectors (Promega, Madison, WI) were used for the promoter reporter assays. The promoter construct of the IL-6 gene was generated by PCR amplification of a 349 bp fragment that was then inserted the KpnI/HindIII fragments of the human IL-6 gene promoter into pGL4.10-Basic vector. The resulting plasmid was named pGL4.10-IL-6 construct. Promoter constructs containing nucleotide substitutions in the sequence motifs of AP1, NF-κB, Sp1 and 5 GR sites were individually generated by PCR amplification with primer pairs spanning the mutant nucleotides according to the protocol of site-directed mutagenesis by overlap extension [18]. The GR cDNA fragment containing NheI and XbaI sites was cloned by overlap extension using RT-PCR and inserted into pCMS-EGFP, a mammalian expression vector. The sequences of primers used in cloning IL-6 promoter and GR cDNA fragments and site-directed mutagenesis are shown in supplemental information.

### Transient transfection

Exponentially growing RAW264.7 cells were seeded at a density of 2.0×10^5^cells per well in 12-well plates. Twenty-four hours later, transient transfection was performed using PolyJet DNA transfection reagent (SignaGen Laboratories, Rockville, MD, USA) according to the manufacture’s protocol at a transfection reagent/DNA ratio of 1:3. Then, plasmid DNA was extracted purification with the UltraClean Endotoxin removal kit (MoBio Lab, Carlsbad, CA) before co-transfection. The plasmid mixtures containing 0.5μg of pGL4.10-IL-6 plasmid and 0.5μg of pGL4.74 [*hRluc*/TK] vector were co-transfected for 24 h. After the medium was changed, LPS (1μg/ml) and/or DEX (10 μM/ml) were added to the plates.

### Dual-luciferase assay

After LPS (1μg/ml) and/or DEX (10μM/ml) treatment for 24 h, the cells were washed with PBS and then lysates were prepared by scraping the cells from plates in the presence of 1× Passive lysis buffer (Promega). Luciferase assays were performed according to Dual-Luciferase Assay System (Promega) and then detected by a Sirius luminometer (Berthold Detection System, Pforzheim, Germany).

### Nuclear protein extraction of human IMR-32 cells

IMR-32 cells were collected from culture dish and centrifuged for 5 minutes at 500 x g. The cell pellets were mixed with 10-fold volume of buffer A (10 mM HEPES (pH 7.9), 1.5 mM MgCl_2_, 10 mM KCl, 0.5 mM DTT, 1 tablet of protease and phosphatase inhibitor (Thermo Scientific, USA)) and incubated on ice for 8 minutes followed by shortly centrifuged for 10 seconds at 12,000 x g at 4 °C. The buffer A-treated cell pellets were then mixed with 2-fold volume of buffer C (20 mM HEPES (pH 7.9), 25% glycerol, 420 mM NaCl, 1.5 mM MgCl_2_, 0.2 mM EDTA, 0.5 mM DTT, 1 tablet of protease and phosphatase inhibitor) and incubated on ice for 16 minutes. After 12,000 x g centrifugation at 4°C, the supernatant containing the nuclear protein of IMR-32 cells were collected and stored at −80°C.

### Gel electrophoretic mobility shift assays (EMSA)

In EMSA, The DNA probes with putative GR-binding sites were prepared by annealing the two complementary biotin-labeled single-strand oligonucleotides as following steps: an equal amount of complementary oligos was mixed in 1x DNA-binding buffer and heated at 94 °C for 2 minutes, and then gradually cooled down to 25 °C. In this study, EMSA was performed by LightShift^®^ Chemiluminescent EMSA Kit (Thermo Scientific, USA) according to the manufacturer’s instructions. To decrease the possibility of non-specificity binding, the nuclear protein from IMR-32 cells were first incubated with 62.5 ng poly(dI-dC)•poly(dI-dC) at 4°C in the 1 x binding buffer for 30 minutes. For competition experiments, excess of unlabeled DNA was added in the binding reaction. Next, the biotin-labeled DNA probes was added and incubated at 4 °C for 30 minutes. The 20 μLof mixtures were run electrophoresis on a 4% native gel at 100 volts, and then transferred onto a 0.45 μM positive-charged nylon membrane (Pall Corporation, USA) at 380 mA in the cold 0.5X TBE buffer (VWR International, USA). The bands on the membrane were visualized through Streptavidin-HRP hybrid followed by enhanced chemiluminescence treatment.

### Chromatin Immunoprecipitation (ChIP) Assay [19]

ChIP assays were performed with an EZ ChIP kit purchased from Upstate Biotechnology (Millipore Corporation Billerica, MA). Briefly, Raw 264.7 cells were fixed with 1% formaldehyde for 10 min at RT for DNA cross linking. The chromatin was sheared by sonication under optimized conditions. ChIP reactions were conducted following the manufacturer’s protocol with the anti-GR, anti-RNA polymerase II, and anti–mouse IgG (Santa Cruz, CA). The IL-6 promoter region was amplified with the primer pair mIL-6 F1 and mIL-6 R3. The polymerase chain reaction (PCR) cycle parameters were: 41 cycles of 95 °C for 20 sec, 58 °C for 30 sec, and 72 °C for 30 sec, and a final extension for 15 min at 72 °C. PCR products were separated by agarose gel electrophoresis.

### Statistical analyses

All values are expressed as means ± SD. Groups were compared using Student’s two-tailed unpaired t-test. A value of *P* < 0.05 was considered to be statistically significant.

## Results

### The LPS-induced morphology changes of RAW264.7 and its IL-6 gene expression reversed by co-treated dexamethasone

To identify the LPS-induced effects of glucocorticoids on IL-6 gene expression, we treated LPS and DEX in RAW264.7 cells and measured the promoter activities and RNA levels of IL-6 gene. We found that the morphology of RAW264.7 cells changed under LPS treatment (Fig 1A). To study the regulation of IL-6 gene, we constructed a 349-bp promoter fragment of IL-6 gene into the pGL4.10-Basic vector, which contains a reporter luciferase gene for measuring promoter activity. We also used RT-PCR to detect IL-6 gene transcript. We found that the promoter activities and mRNA levels were induced by LPS and reversed by DEX (Figs 1B and C). However, treatment of DEX alone did not change the cellular morphology, promoter activities and mRNA levels of IL-6 gene. These results indicate that glucocorticoids modulate LPS-induced morphological change and IL-6 gene expression.

**Figure 1.**
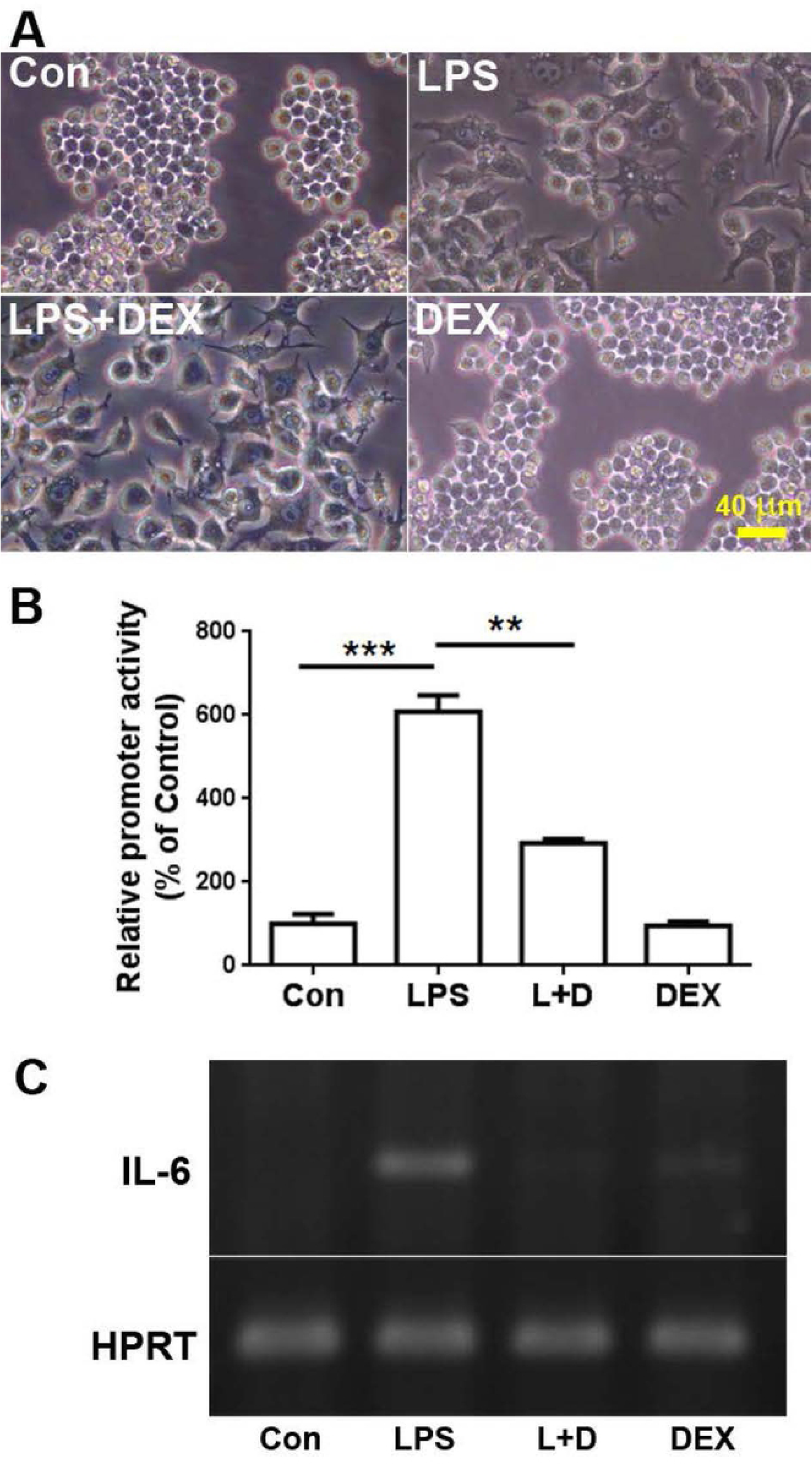
Dexamethasone downregulates LPS-induced macrophage activation and IL-6 gene expression. (A) Morphological changes were induced by LPS and dexamethasone. RAW264.7 cells were treated with 1 μg/ml LPS and 10μM dexamethasone for 24 h. (B) Dexamethasone (DEX) mediated the LPS-induced IL-6 promoter activities. (C) The mRNA levels of IL-6 were induced by LPS and revered by adding DEX. One representative image obtained from one of three individual experiments is shown.

### Different changes of promoter activities in mutant putative sites on IL-6 promoter by using site-directed mutagenesis method

To predict which proteins bind to IL-6 promoter to mediate the gene expression, we searched the transcription factor binding sites on IL-6 promoter region using bioinformatics tools and found several putative sites, such as AP-1, NF-κB, Sp1 and 5 GR sites. The sequences of these sites show high similarity among different species (Supplemental material, Fig S1). To clarify which sites are important for regulating IL-6 gene expression, we used the site-directed mutagenesis to construct the mutant IL-6 promoter-reporter vectors and transfected theses constructs to RAW264.7 cells. Mutation of AP-1 site decreased the basal and effects of LPS-induced and addition of DEX in IL-6 promoter activities (Fig 2A). Importantly, mutation of NF-κB site dramatically decreased the promoter activities (Fig 2B). There are two Sp1 sites on IL-6 promoter, denoted Sp1-1 and Sp1-2. Mutation of Sp1-2 but not Sp1-1 sites decreased IL-6 promoter activity, suggesting these two sites exert differential function in IL-6 expression in this cell line (Fig 2C). These results suggest that AP-1 and NFκB sites are important for IL-6 express in RAW264.7 cells but only one Sp1 site is partially involved in this process.

**Figure 2.**
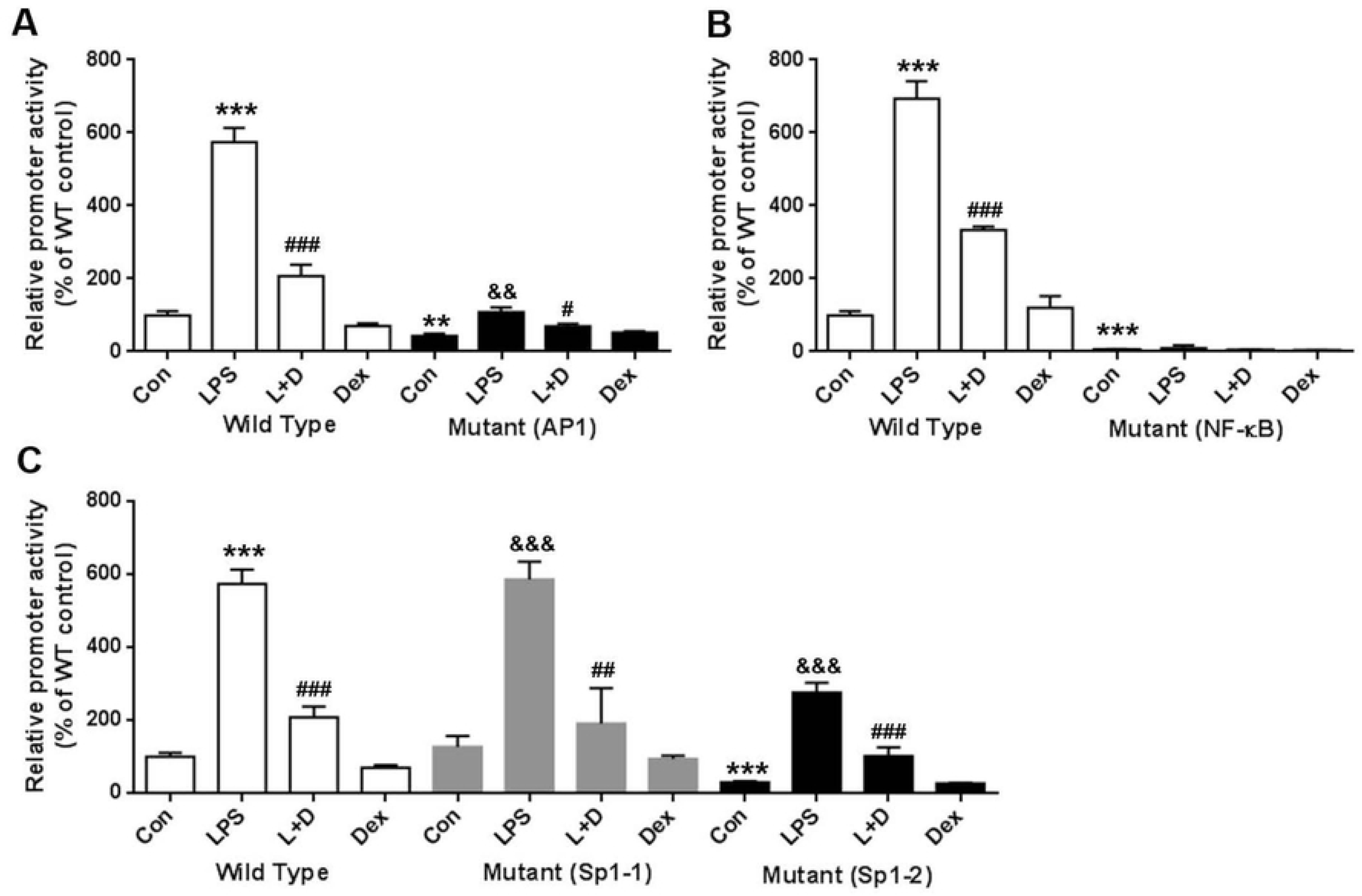
Effects of mutation of the putative transcriptional factor binding sites. (A) Promoter activities of mutant AP-1 sites were significantly decreased compared with wild-type controls. (B) Promoter activities of mutant NF-kB sites were more significantly decreased than wild controls. (C) Promoter activities of mutant Sp1-2 sites showed partially decreased change compared with wild-type controls; however, there were no significant change between mutant Sp1-1 sites and wild-type controls.

### The basal and inducible changes of IL-6 promoter activities in mutant five GR binding sites and the reversed effect of dexamethasone treatment

There are five putative GR binding sites on IL-6 promoter. We transfected the mutant promoter constructs of GR sites to detect which sites are important for the IL-6 promoter activities and the results showed that mutation the GR2 and 3 sites decreased the basal and LPS-induced promoter activities. Furthermore, the DEX-reversed effects were also altered. However, mutation of the GR1, 4 and 5 sites did not affect the effects of LPS and DEX treatments (Fig 3). These results suggest that GR2 and GR3 are important for IL-6 promoter activities.

**Figure 3.**
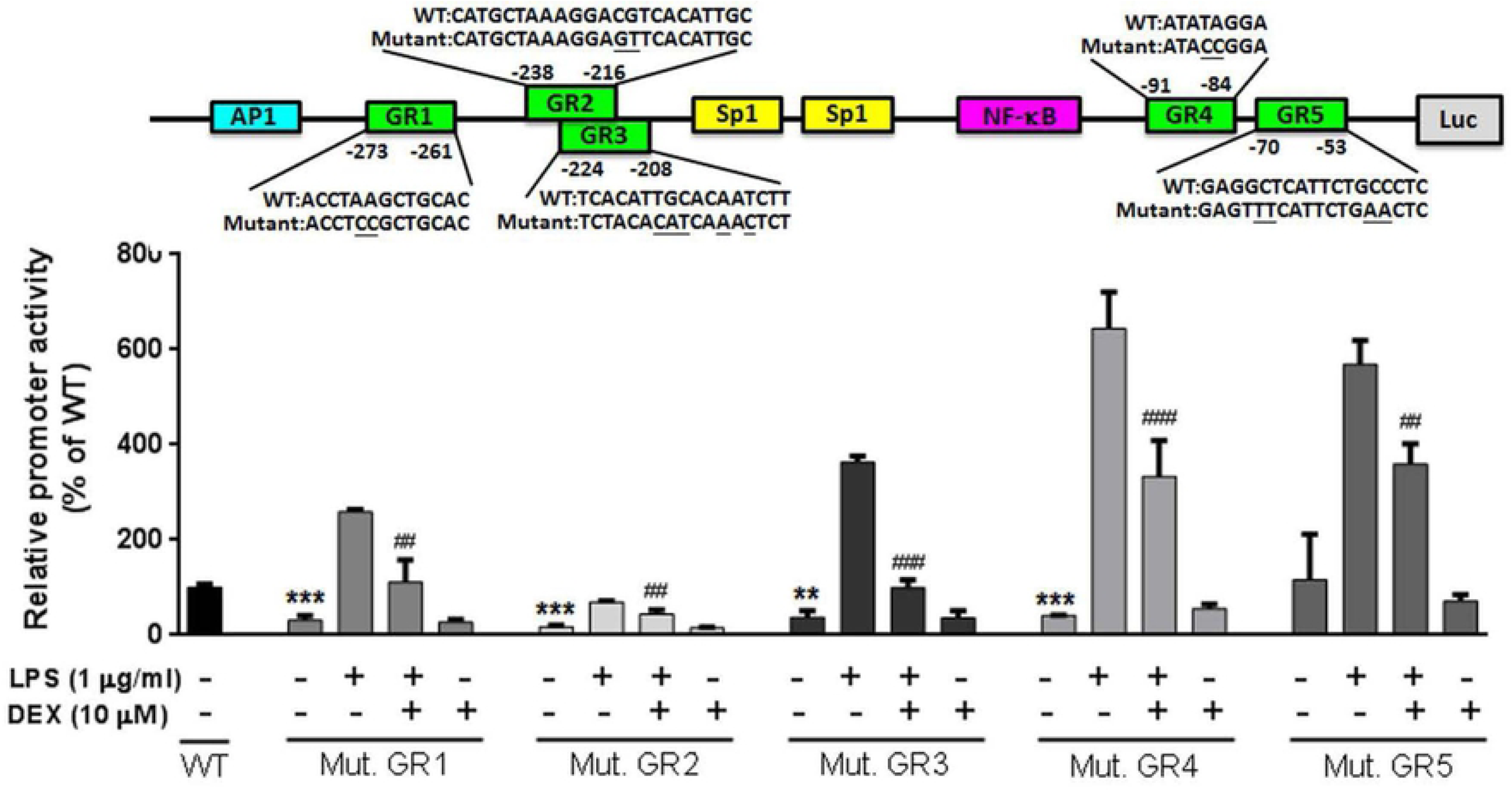
Mutation of putative GR2/3 binding sites reduces the basal promoter activity of IL-6 gene. Upper shows the mutant sequences of five GR binding sites. Lower shows the promoter activities of mutant GR sites. Among them, mutant GR2 binding site showed a significant decrease compared with other mutant binding sites.

### The changes of IL-6 promoter activities and gene expression under over-expression of GR

To understand the role of GR in regulating the expression of IL-6, we over-expressed GR in RAW264.7cells by transfecting various doses of GR cDNA constructs and found the IL-6 promoter activities were increased in a dose dependent manner (Fig 4A). To identify if GR binds to IL-6 promoter, we used ChIP and EMSA to investigate the protein-DNA binding in vitro and in vivo, respectively. In the ChIP assay, GR can bind to IL-6 promoter indeed (Fig 4B). In the EMSA assay, the sequence analysis shows that GR2 and 3 overlap partially, so we used one probe containing these two sites for EMSA. The EMSA showed that the probe of GR 2/3 site but not others presents shifted bands. Furthermore, we used GR binding motif of elastin promoter as cold probes for competition assay and found the shifted bands were eliminated (Fig 4C), suggesting that GR binds to GR2/3 sites. However, there are no sifted bands in EMSA using the probes containing GR1, 4 and 5 sites. According to these results, we confirmed that GR binds to GR2/3 sites but not others on the IL-6 promoter region.

**Figure 4.**
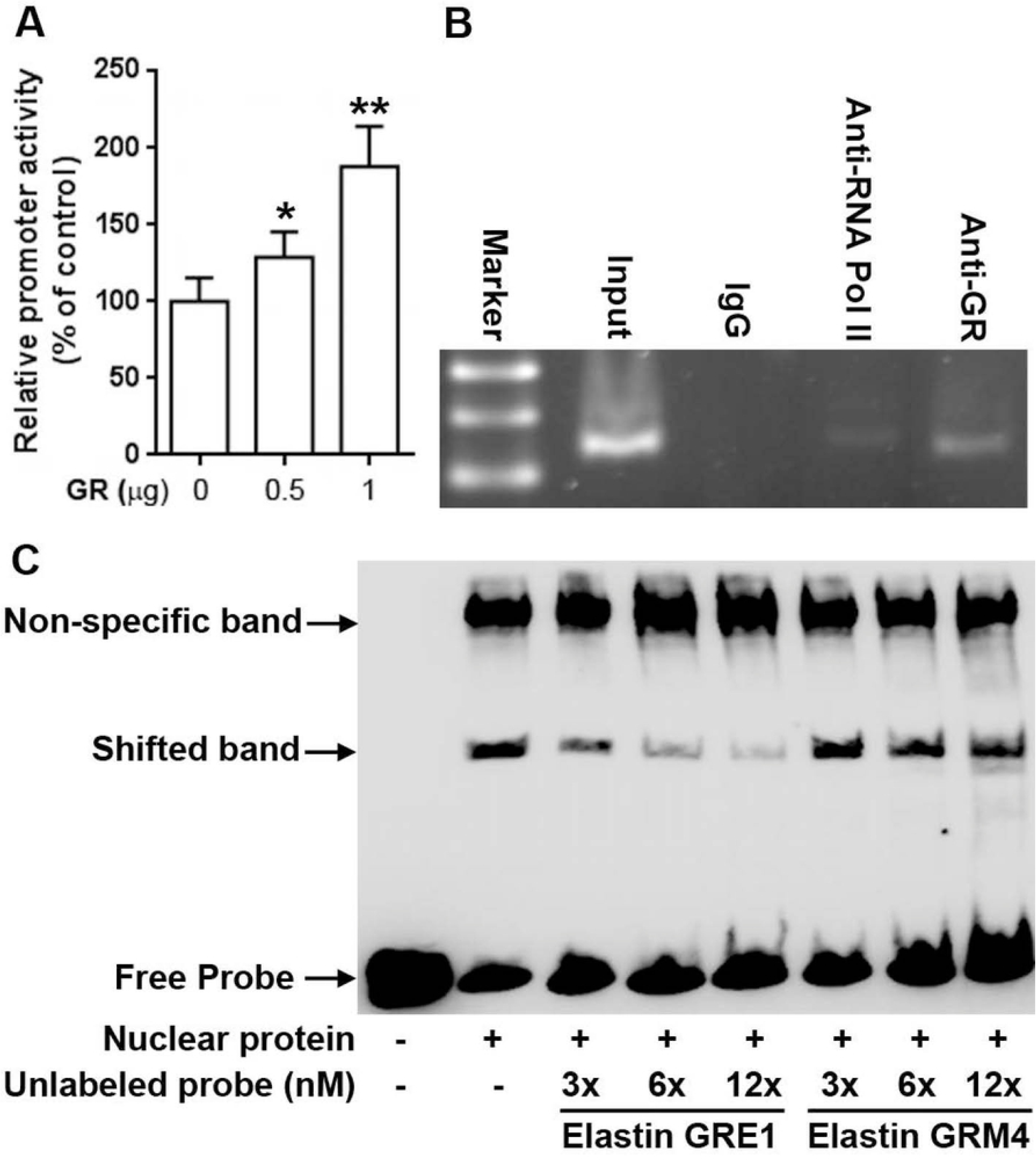
GR directly binds to IL-6 promoter and contributes to its activity. (A) Over-expression of GR increased IL-6 promoter activities in a dose-dependent manner. (B) GR binds to IL-6 promoter region in vivo. Anti-mouse IgG and anti-RNA polymerase II antibodies were used for the negative and positive controls in ChIP assay, respectively. (C) Competition assay presents the GR binding on GR 2/3 sites of IL-6 promoter. The fragments containing wild type and mutant GR binding motifs were used as competitors.

### The proposed schematic model for dexamethasone regulates LPS-stimulated IL-6 expression via GRs

We proposed the schematic model for DEX negative regulates LPS-stimulated IL-6 expression in RAW 264.7 cells (Fig 5). The LPS stimulates nuclear IL-6 promoter activity through NF-κB pathway and the counter-regulation of NF-κB by the GR was proposed on IL-6 promoter.

**Figure 5.**
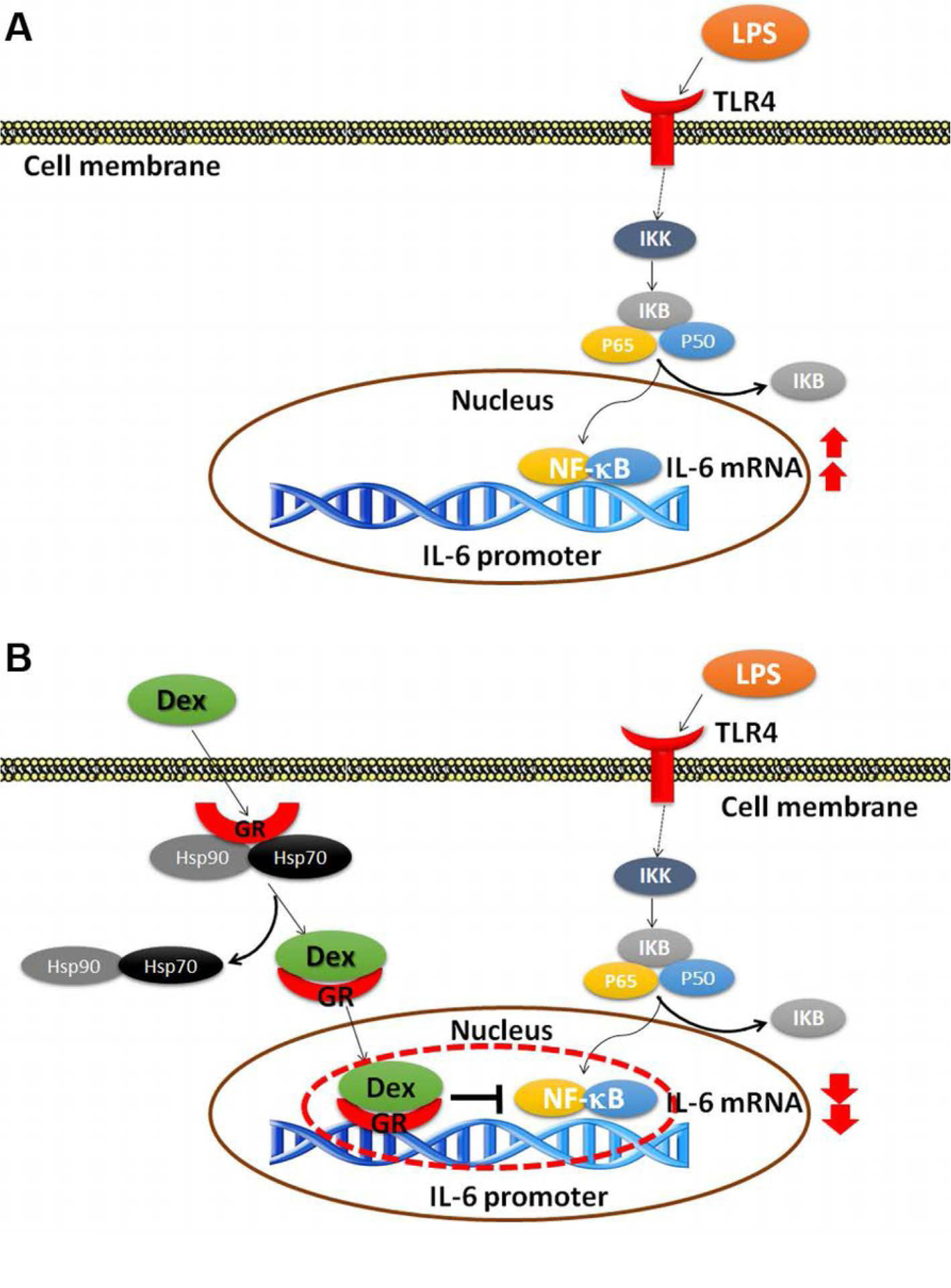
The schematic model for regulation of dexamethasone in LPS-stimulated IL-6 expression. (A) LPS stimulates IL-6 promoter activity thru NF-κB pathway in RAW 264.7 cells. (B) A proposed mechanism for counter-regulation of NF-κB by dexamethasone and GR on IL-6 promoter.

## Discussion

Our studies show that the pretreatment of Raw 264.7 cells with dexamethasone at different concentrations could significantly decrease the LPS-induced mRNAs expression of Il-6 in a dose dependent manner. It suggested that the protective role of synthetic dexamethasone effects on LPS-induced inflammation. Our results consistent with the findings of Yamazaki et al., who reported that the GCs inhibit the synthesis of interleukin (IL)-1, TNF-α, IL-1β, IL-6, MMP-I, and COX-2 mRNAs expression in SW982 cells [20].

The molecular mechanism of GCs through interaction with their receptor GRs to regulate the inflammation responses has been well-studied. GCs bind to GRs and are transported into the nucleus subsequently to form a dimer and also combinations of transcriptional complexes, such as AP-1 and NF-κB, either in transrepression or in transactivation, which are responsible for anti-inflammatory action of GCs [21,22]. The results from previous studies suggest that the treatment of naïve monocytes with fluticasone or dexamethasone, did not cause a global suppression of the activated monocytic functions instead of induction of cellular differentiation with an anti-inflammatory phenotype [23,24]. Recently, growing evidences show that GCs could efficiently inhibit these processes by down-regulating pro-inflammatory mediators from macrophages and monocytes and their migration toward inflammatory stimuli. GCs remove endo- and exogenous danger signals by an increased phagocytic capacity, and limit T-cell activation [25].

Our results showed that LPS stimulation resulted in up-regulated expressions of inflammatory transcription factors such as NF-κB [26], AP-1 [27] and Sp1-2 [28]. LPS also resulted in the increases of IL-6 and its promoter activity. The increase of LPS-stimulated IL-6 promoter activities could be suppressed by DEX, esp. in NF-κB and AP-1, Sp1-2, but not in Sp1-1. These results are consistent with previous well-known studies [29,30]. It has been shown in the previous study that two Sp1 sites on IL-6 promoter and Sp1 binds to this region indeed in the human monocytes [31]. However, there is no report to elucidate which Sp1 site are important for IL-6 expression. We first discovered that only the Sp1-2 site is involved in IL-6 expression. Taken together, the transcription factors NF-κB, AP-1, Sp1-2 bind to IL-6 promoter and play important roles in basal and in inducible expression of the murine IL-6 gene.

In our study, we have found five putative GRs binding sites on IL-6 promoter and DEX could decrease the promoter activities of IL-6 GR binding sites. The promoter activity of IL-6 after LPS stimulation and the construct sequence, showed a significant increase of LPS-inducible promoter activity. Among them, GR2/3 seemed to be an important role in both basal and inducible promoter activities in LPS-induced inflammation. Comparing to GR1, GR4, GR5, they did not show the similar changes of LPS-inducible or basal promoter activities. Although previous study shows that recombinant GR binds to IL-6 promoter regions respect to GR1-5 elements in vitro [10], there were no significant shifted bands using the probes of GR1, GR4, GR5 for EMSA and neither changes of the promoter activities being found after mutation of these three sites (data not shown). It has been proposed that GR interacts with transcription factors, such as AP-1and NF-κB, to mediate the downstream proinflammatory genes, including IL-6 [32]. The exact targeted regulation of GR2/3 on IL-6 and the possible mechanism related to transcriptional synergism with GRE remain unknown. Whether GR2/3 plays a novel negative GRE [33], directly binding by GRα, the pivotal subunit of GR, which could involve the interactions between GRα and AP-1and NF-κB, needs further studies to identify.

Our results provide the possible mechanism of controversial steroids should be administrated in hemodynamic unstable septic patient whether in low- or high doses. The GR2/3 binding site might provide the important selection of candidate therapeutics in vulnerable septic patients in the future.

## Acknowledgements

not applicable

## Funding

This work was supported in part by NCKUH, IRB No: B-ER-102-121 (2013) and B-ER-103-419 (2014).

## Competing interests

None of the authors have financial and non-financial competing interests.

## Authors’ Contributions

Chang WT and Chuang CC designed the study. Hong MY and Hwang CY prepared the data. Chen CL helped the statistical analyses. Chang WT and Tsai CC performed the experiment. Tsai CC and Chang WT drafted the initial manuscript. Chang WT and Chuang CC performed the final review and editing. All authors reviewed and approved the final version of this manuscript.

## References

1. Chousterman BG, Swirski FK, Weber GF. Cytokine storm and sepsis disease pathogenesis. Semin Immunopathol. 2017;39:517–28. doi: 10.1007/s00281-017-0639-8. PMID: 28555385.

2. Cohen J, Pretorius CJ, Ungerer JP, Cardinal J, Blumenthal A, Presneill J, et al. Glucocorticoid sensitivity is highly variable in critically ill patients with septic shock and is associated with disease severity. Crit Care Med. 2016;44:1034–41. doi: 10.1097/ccm.0000000000001633. PMID: 26963327.

3. Fang F, Zhang Y, Tang J, Lunsford LD, Li T, Tang R, et al. Association of corticosteroid treatment with outcomes in adult patients with sepsis: A systematic review and meta-analysis. JAMA Intern Med. 2019;179:213–23. doi: 10.1001/jamainternmed.2018.5849. PMID: 30575845.

4. Rygard SL, Butler E, Granholm A, Moller MH, Cohen J, Finfer S, et al. Low-dose corticosteroids for adult patients with septic shock: a systematic review with meta-analysis and trial sequential analysis. Intensive Care Med. 2018;44:1003–16. doi: 10.1007/s00134-018-5197-6. PMID: 29761216.

5. Wang C, Sun J, Zheng J, Guo L, Ma H, Zhang Y, et al. Low-dose hydrocortisone therapy attenuates septic shock in adult patients but does not reduce 28-day mortality: a meta-analysis of randomized controlled trials. Anesth Analg. 2014;118:346–57. doi: 10.1213/ane.0000000000000050. PMID: 24445635.

6. Stahn C, Lowenberg M, Hommes DW, Buttgereit F. Molecular mechanisms of glucocorticoid action and selective glucocorticoid receptor agonists. Mol Cell Endocrinol. 2007;275:71–8. doi: 10.1016/j.mce.2007.05.019. PMID: 17630118.

7. Baschant U, Tuckermann J. The role of the glucocorticoid receptor in inflammation and immunity. J Steroid Biochem Mol Biol. 2010;120:69–75. doi: 10.1016/j.jsbmb.2010.03.058. PMID: 20346397.

8. Patel GP, Balk RA. Systemic steroids in severe sepsis and septic shock. Am J Respir Crit Care Med. 2012;185:133–9. doi: 10.1164/rccm.201011-1897CI. PMID: 21680949.

9. Li CC, Munitic I, Mittelstadt PR, Castro E, Ashwell JD. Suppression of dendritic cell-derived IL-12 by endogenous glucocorticoids is protective in LPS-induced sepsis. PLoS Biol. 2015;13:e1002269. doi: 10.1371/journal.pbio.1002269. PMID: 26440998.

10. Ray A, LaForge KS, Sehgal PB. On the mechanism for efficient repression of the interleukin-6 promoter by glucocorticoids: enhancer, TATA box, and RNA start site (Inr motif) occlusion. Mol Cell Biol. 1990;10:5736–46. doi: 10.1128/mcb.10.11.5736. PMID: 2233715.

11. Waage A, Slupphaug G, Shalaby R. Glucocorticoids inhibit the production of IL6 from monocytes, endothelial cells and fibroblasts. Eur J Immunol. 1990;20:2439–43. doi: 10.1002/eji.1830201112. PMID: 2253684.

12. Beck IM, Vanden Berghe W, Vermeulen L, Yamamoto KR, Haegeman G, De Bosscher K. Crosstalk in inflammation: the interplay of glucocorticoid receptor-based mechanisms and kinases and phosphatases. Endocr Rev. 2009;30:830–82. doi: 10.1210/er.2009-0013. PMID: 19890091.

13. Ng HP, Jennings S, Wang J, Molina PE, Nelson S, Wang G. Noncanonical glucocorticoid receptor transactivation of gilz by alcohol suppresses cell inflammatory response. Front Immunol. 2017;8:661. doi: 10.3389/fimmu.2017.00661. PMID: 28638383.

14. Xavier AM, Anunciato AK, Rosenstock TR, Glezer I. Gene expression control by glucocorticoid receptors during innate immune responses. Front Endocrinol (Lausanne). 2016;7:31. doi: 10.3389/fendo.2016.00031. PMID: 27148162.

15. Roman-Blas JA, Jimenez SA. NF-kappaB as a potential therapeutic target in osteoarthritis and rheumatoid arthritis. Osteoarthritis Cartilage. 2006;14:839–48. doi: 10.1016/j.joca.2006.04.008. PMID: 16730463.

16. Patil RH, Naveen Kumar M, Kiran Kumar KM, Nagesh R, Kavya K, Babu RL, et al. Dexamethasone inhibits inflammatory response via down regulation of AP-1 transcription factor in human lung epithelial cells. Gene. 2018;645:85–94. doi: 10.1016/j.gene.2017.12.024. PMID: 29248584.

17. Chuang CC, Chuang YC, Chang WT, Chen CC, Hor LI, Huang AM, et al. Macrophage migration inhibitory factor regulates interleukin-6 production by facilitating nuclear factor-kappa B activation during Vibrio vulnificus infection. BMC Immunol. 2010;11:50. doi: 10.1186/1471-2172-11-50. PMID: 20939898.

18. Aiyar A, Xiang Y, Leis J. Site-directed mutagenesis using overlap extension PCR. Methods Mol Biol. 1996;57:177–91. doi: 10.1385/0-89603-332-5:177. PMID: 8850005.

19. Wang JL, Chang WT, Tong CW, Kohno K, Huang AM. Human synapsin I mediates the function of nuclear respiratory factor 1 in neurite outgrowth in neuroblastoma IMR-32 cells. J Neurosci Res. 2009;87:2255–63. doi: 10.1002/jnr.22059. PMID: 19301426.

20. Yamazaki T, Tukiyama T, Tokiwa T. Effect of dexamethasone on binding activity of transcription factors nuclear factor-kappaB and activator protein-1 in SW982 human synovial sarcoma cells. In Vitro Cell Dev Biol Anim. 2005;41:80–2. doi: 10.1290/0502011.1. PMID: 16029077.

21. Newton R, Holden NS. Separating transrepression and transactivation: a distressing divorce for the glucocorticoid receptor? Mol Pharmacol. 2007;72:799–809. doi: 10.1124/mol.107.038794. PMID: 17622575.

22. Segard-Maurel I, Rajkowski K, Jibard N, Schweizer-Groyer G, Baulieu EE, Cadepond F. Glucocorticosteroid receptor dimerization investigated by analysis of receptor binding to glucocorticosteroid responsive elements using a monomer-dimer equilibrium model. Biochemistry. 1996;35:1634–42. doi: 10.1021/bi951369h. PMID: 8634295.

23. Ehrchen J, Steinmuller L, Barczyk K, Tenbrock K, Nacken W, Eisenacher M, et al. Glucocorticoids induce differentiation of a specifically activated, anti-inflammatory subtype of human monocytes. Blood. 2007;109:1265–74. doi: 10.1182/blood-2006-02-001115. PMID: 17018861.

24. Varga G, Ehrchen J, Tsianakas A, Tenbrock K, Rattenholl A, Seeliger S, et al. Glucocorticoids induce an activated, anti-inflammatory monocyte subset in mice that resembles myeloid-derived suppressor cells. J Leukoc Biol. 2008;84:644–50. doi: 10.1189/jlb.1107768. PMID: 18611985.

25. Ehrchen JM, Roth J, Barczyk-Kahlert K. More than suppression: glucocorticoid action on monocytes and macrophages. Front Immunol. 2019;10:2028. doi: 10.3389/fimmu.2019.02028. PMID: 31507614.

26. Liu T, Zhang L, Joo D, Sun SC. NF-kappaB signaling in inflammation. Signal Transduct Target Ther. 2017;2. doi: 10.1038/sigtrans.2017.23. PMID: 29158945.

27. Cahill CM, Zhu W, Oziolor E, Yang YJ, Tam B, Rajanala S, et al. Differential expression of the activator protein 1 transcription factor regulates interleukin-1ss induction of interleukin 6 in the developing enterocyte. PLoS One. 2016;11:e0145184. doi: 10.1371/journal.pone.0145184. PMID: 26799482.

28. Ye X, Liu H, Gong YS, Liu SF. LPS Down-regulates specificity protein 1 activity by activating NF-kappaB pathway in endotoxemic mice. PLoS One. 2015;10:e0130317. doi: 10.1371/journal.pone.0130317. PMID: 26103469.

29. Steer JH, Kroeger KM, Abraham LJ, Joyce DA. Glucocorticoids suppress tumor necrosis factor-alpha expression by human monocytic THP-1 cells by suppressing transactivation through adjacent NF-kappa B and c-Jun-activating transcription factor-2 binding sites in the promoter. J Biol Chem. 2000;275:18432–40. doi: 10.1074/jbc.M906304199. PMID: 10748079.

30. Saiki P, Nakajima Y, Van Griensven L, Miyazaki K. Real-time monitoring of IL-6 and IL-10 reporter expression for anti-inflammation activity in live RAW 264.7cells. Biochem Biophys Res Commun. 2018;505:885–90. doi: 10.1016/j.bbrc.2018.09.173. PMID: 30301531.

31. Sanceau J, Kaisho T, Hirano T, Wietzerbin J. Triggering of the human interleukin-6 gene by interferon-gamma and tumor necrosis factor-alpha in monocytic cells involves cooperation between interferon regulatory factor-1, NF kappa B, and Sp1 transcription factors. J Biol Chem. 1995;270:27920–31. doi: 10.1074/jbc.270.46.27920. PMID: 7499267

32. De Bosscher K, Vanden Berghe W, Haegeman G. The interplay between the glucocorticoid receptor and nuclear factor-kappaB or activator protein-1: molecular mechanisms for gene repression. Endocr Rev. 2003;24:488–522. doi: 10.1210/er.2002-0006. PMID: 12920152.

33. Luo Y, Zheng SG. Hall of fame among pro-inflammatory cytokines: interleukin-6 gene and its transcriptional regulation mechanisms. Front Immunol. 2016;7:604. doi: 10.3389/fimmu.2016.00604. PMID: 28066415

